# *Toll-like receptor 3* orchestrates a conserved mechanism of heart regeneration

**DOI:** 10.1101/2024.04.12.589327

**Authors:** Manuel Fiegl, Bernhard Roehrs, Dominik Hau, Sophia Mair, Philipp Niederwanger, Dominik Regele, Lorenzo Merotto, Sonja Toechterle, Clemens Engler, Michael Graber, Elke Kirchmair, Veronika Niedrist, Lynn Muller, Leo Poelzl, Jakob Hirsch, Felix Naegele, Isidor Happacher, Simon Oberegger, Nikolaos Bonaros, Michael Grimm, Ivan Tancevski, Francesca Finotello, Dirk Meyer, Johannes Holfeld, Can Gollmann-Tepekoeylue

## Abstract

The human’s heart responds to tissue damage with persistent fibrotic scarring. Unlike humans, zebrafish can repair cardiac injury and re-grow heart tissue throughout life. Recently, *Toll-like receptor 3* (*Tlr3*) was identified as an important mediator of cardiac regeneration in neonatal mice. However, no functional analysis of *tlr3* knock-out mutant zebrafish in respect to cardiac regeneration has yet been performed.

We hypothesize that TLR3 signalling plays a central, conserved role in driving cardiac regeneration upon injury. Therefore, we focused on *tlr3* mediated cardiac regeneration in zebrafish, ultimately discovering an evolutionary conserved mechanism of heart repair. Using histological, behavioural, and RNA-Sequencing analysis, we uncovered a conserved mechanism of *tlr3* mediated cardiac repair after myocardial injury.

Upon myocardial cryoinjury subjection, survival is decreased in *tlr3^-/-^* fish as compared to wildtype controls. *Tlr3^-/-^* zebrafish fail to recruit immune cells to the injured ventricle, resulting in impaired DNA repair and transcriptional reprogramming of cardiomyocytes.

Mechanistically, we uncover an evolutionary conserved mechanism of *tlr3* activation in fibroblasts promoting monocyte migration towards an injured ventricular area. Our data reveal *tlr3* as a novel therapeutic target to promote cardiac regeneration.

Every experiment including human participants has been approved by the ethics committee of the Medical University of Innsbruck (Ref. Nr.: 1262/2023). All experiments including the use of laboratory animals have been approved by the federal ministry of education, science, and research of Austria (Ref. Nr.: 2020-0.345.504).

## Introduction

In patients with ischaemic heart disease (IHD), blood flow in the heart muscle is reduced, resulting in oxygen deprivation and tissue loss. Given the restricted regenerative capacity of human cardiac tissue upon injury, IHD represents the leading cause of death in the Western world^1^. Adult mammalian cardiac tissue was long considered as post-mitotic, with a limited ability to proliferate and replace damaged tissue^2^. However, research in neonatal mice has revealed their ability to regenerate cardiac injuries until seven days after birth^3^. Furthermore, induction of DNA-damage repair mechanisms improves cardiac function and cardiomyocyte proliferation even in adult mice, resulting in reduced scarring after myocardial infarction^4^. These findings suggest that a certain cardiac regenerative capacity might as well be conserved in humans, and that the latent ability to regenerate heart tissue is actively suppressed during maturation^5^. Therefore, investigation of species able to retain their cardiac regenerative capacity throughout life has recently gained interest.

The zebrafish is among the most relevant animal models to study regenerative biology due to its impressive capacity of organ regeneration, including the heart^6–9^. Upon cardiac injury, massive cell death events trigger an initial pro-inflammatory phase. Attracted by damage-associated molecular patterns (DAMPs) and cytokines released from damaged tissue, neutrophils and macrophages infiltrate the affected area immediately after injury, clearing debris and dead cells while also mediating extracellular matrix (ECM) remodelling^2,10,11^.

Canonically, macrophages exist in a range of diverse activation states orchestrated by specific cytokine signatures^12,13^. Different macrophage subtypes exhibit bifunctional roles during myocardial regeneration of zebrafish, being involved in collagen deposition first, while later contributing to resolving the initially formed fibrotic scar and replacing it with functional cardiac tissue^14^. Collagens are the most abundant protein constituent of the ECM with 59 predicted collagen genes^15,16^. Besides their main function in stabilizing the ventricle upon injury, collagens and other cardiac ECM proteins play a vital role in mechanical and biochemical interactions influencing cell behaviour and function. Cardiac injury causes dynamic changes in the ECM that contribute to regulation of inflammation, repair and potentially mediate cardiac remodelling^17^. Immune cells clear the injured area from dead cardiac tissue, while injury-border zone located cardiomyocytes undergo limited dedifferentiation characterized by detachment from one another and cell cycle re-entry. Dedifferentiated cardiomyocytes then start proliferating and ultimately replace the fibrotic scar, resulting in almost complete tissue regeneration^2,18^.

*Toll-like receptor 3* (TLR3), a central element of the innate immune system, was recently identified as an important mediator of cardiac regeneration in neonatal mice. *Tlr3* deficiency impaired cardiac regeneration after myocardial infarction (MI) through inhibition of cardiomyocyte cell cycle re-entry and proliferation^19^. Moreover, TLR3 is implicated in regeneration of other tissues, being able to promote spinal cord repair upon injury in both mice and zebrafish^20^. In adult mice, Tlr3 activation improves myocardial function through modulation of inflammatory response^21^. TLR3 canonically binds double-stranded RNA (dsRNA), inducing the expression of proinflammatory cytokines such as type I Interferons (IFN) and *Interleukin-6* (IL6), mediated by their transcription factors *Interferon regulatory factor 3* (IRF3) and *Nuclear factor k-light-chain-enhancer of activated B cells* (NF-κB), respectively^21^. IL6 and NF-κB signalling are both essential for zebrafish cardiac regeneration, blocking cardiomyocyte proliferation upon functional inhibition^22,23^. However, no functional analysis of *tlr3* knock-out mutant zebrafish in respect to cardiac regeneration has yet been performed.

In this project, we subjected *tlr3* knock-out mutant zebrafish to myocardial infarction through cryoinjury. Survival of *tlr3* mutants was markedly impaired compared to wild types, while surviving mutant fish failed to resolve their initially formed scar. Mutant injured hearts displayed decreased neutrophil and macrophage recruitment, resulting in increased heterogeneity of collagen gene expression, while DNA-repair & muscle stem cell signatures were decreased. Ultimately, we postulate a potential conserved mechanism of *tlr3* activation in fibroblasts to mediate immune cell migration towards the injured ventricular area and drive monocyte maturation through the release of cytokines such as *Monocyte Chemotactic Protein 1, 2, 3 (MCP-1, MCP-2, MCP-3) & Colony Stimulating Factor 1* (*CSF-1*). Hence, we demonstrate the crucial role of *tlr3* signalling in cardiac regeneration in a previously undescribed mutant model and reveal a novel target to potentially induce cardiac regeneration in patients with ischemic cardiomyopathy.

## Methods

### Zebrafish handling

All zebrafish (*Danio rerio*, strain: Tüb/AB) husbandry was performed according to standard procedures^36^, and all experiments have been approved by the federal ministry of education, science, and research of Austria (Ref. Nr.: 2020-0.345.504).

### Zebrafish lines

The following zebrafish lines were used in this study: Wildtype (strain: Tüb/AB), abbreviated as WT; *tlr3* homozygous knock out mutants generated, abbreviated as either *tlr3* mutant or *tlr3^-^/-* (ZFIN ID: ml111).

### Generation of zebrafish transgenic and mutant lines

Mutant zebrafish lines were created in previous works by Röhrs^37^ (ZFIN ID: ml111). CRISPR- Cas9 genome editing technology was used to generate *tlr3* knock-out mutants. The following guide RNA (gRNA) sequences (5′-3′) were used: CACTGGATGTATCTCACACC, GGGCATGGGCATCAACAAGT. gRNAs together with Cas9 protein (TrueCut™ Cas9 Protein v2, Thermo Fisher) was injected into zebrafish embryos at the zygotic/one-cell stage. Corresponding adult fish were outcrossed with WT zebrafish. Mutant alleles were identified by genotyping through PCR, gel electrophoresis and DNA-sequencing. Primer sequences used for genotyping and sequencing are as following: 5’-GCACTACAAATGCACGCAAG-3’ (*tlr3* fwd, also used for sequencing); 5’-CACACCAAACGTAGCCCTTT-3’ (*tlr3* rvs); 5’- ACCATATTCCAGCTGAGCCT-3’ (*tlr3* WT rvs). Heterozygous mutants were in-crossed to generate homozygous mutants. For genotyping, adult zebrafish were anesthetised part of the upper caudal fin isolated with surgical scalpels WT. Mutant alleles were identified through PCR amplifying respective amplicon lengths at 519bp and 700bp.

### Myocardial cryoinjury

The myocardial cryoinjury model was performed according to Chablais et al^38^ with minor adjustments. The accessibility to the ventricular apex was optimized by squeezing the fish posterior to the incision and the incubation time of cryoinjury reduced from 14 to 10 seconds. **Survival analysis**

WT and *tlr3* mutant zebrafish were held in system water upon subjecting them to either cryoinjury or sham treatment. Potential death occurrences were noted at 6hpi, 1dpi, 4dpi and 10dpi. The number of individual fish included in the survival analysis were as following: WT Sham = 22; WT cryoinjured = 127; *tlr3^-/-^* Sham = 26; *tlr3^-/-^* cryoinjured = 87. Probability of survival was assessed through Kaplan-Meier estimate and subsequent log-rank test by Mantel-Cox using GraphPad Prism (v8.4.3) survival analysis tools. Differences in survival were assessed as significant with log-rank test values < 5 x 10^-2^.

### Mobility Assay and Distance to Surface analysis

Movement speed & travelling distance was measured to evaluate mobility behaviour of *tlr3* mutant and WT fish after being subjected to cryoinjury. Untreated, sham treated and cryoinjured fish were kept in mini swarms consisting of 6 – 9 individual fish. At 6hpi, 1dpi, 4dpi and 14dpi, swarms were filmed for a minute from top view, during daytime. Travelling distance per minute in the XY-axis was evaluated using Kinovea software. To evaluate DTS upon injury, *tlr3* mutant and WT fish were subjected to cryoinjury and kept in separate tanks for 4 weeks. For every group, mini swarms consisting of 8 – 11 fish were kept in each tank. Frontal images of the tanks were randomly taken 3 – 5 times per week, during daytime. Average DTS was measured using Fiji (National Institutes of Health, Bethesda, Maryland, USA), calculating the mean DTS of all fish in the tank for daily values and the mean of all daily mean DTS per week for weekly values. Distribution of zebrafish in the tank was assessed using the videos taken while assessing the travelling distance per minute and fish were tracked by hand using the “Manual Tracking” ImageJ plugin.

### Heart collection

Isolation of zebrafish hearts were performed as in Chablais et al^38^.

### Histological analysis & imaging

Isolated zebrafish ventricles connected to the BA, were fixed in 4% Paraformaldehyde (PFA) overnight. Fixed tissue was then dehydrated and infiltrated with Paraffin using an automatic tissue infiltration machine (Leica, TP1020) as follows: 1. Ethanol (70%) / 90min; 2. Ethanol (80%) / 90min; 3. Ethanol (96%) / 90min; 4. Ethanol (100%) / 60min; 5. Ethanol (100%) / 60min; 6. Ethanol (100%) / 60min; 7. Xylene / 90min; 8. Xylene / 90min; 9. Paraffin / 240min; 10. Paraffin / 240min. Fixed ventricles were embedded in embedding cassettes (Carl Roth, 7x7mm) using a Medite TES99 pouring station filled with paraffin wax (Histosec Pastilles, Sigma-Aldrich), longitudinally oriented. Using a Microtome (Leica RM2135), 5µm thick tissue sections were sliced from the paraffin block. Tissue ribbons were transferred to a warm water bath (40°C) and then scooped up onto a charged glass slide (SuperFrost Plus, Thermo Scientific). To obtain 6 replicates for every heart, 6 slides were prepared. About 8 sections per slide were obtained. Tissue sections were dried at 65°C for 2h and then stored at room temperature. For Acid-Fuchsin-Orange-G (AFOG) staining, paraffin sections were deparaffinated in Xylene for 5’ twice and rehydrated in a descending alcohol series (Isopropanol; 96%, 80%, 70%, 60%, 0% (diH_2_O)) for 5’ each. Sections were then fixed in Bouin’s solution (Sigma-Aldrich) for 2 hours at 60°C and stained according to the manufacturer’s instructions, except for skipping the iron-haematoxylin incubation step (Färbekit AFOG / SFOG nach MALLORY & CASON, Morphisto). Lastly, stained sections were embedded in Entellan rapid mounting medium (Sigma-Aldrich) and covered with 24x55mm coverslips (R. Langenbrinck). Imaging was performed using a Zeiss Axioplan 2 microscope and a Zeiss AxioCam HRc digital microscope camera connected to Zeiss imaging software (Zeiss, Oberkochen, Germany).

### Fibrotic scar clearance analysis

WT hearts (n = 3 – 6 per time point) were collected at 4dpi, 30dpi and 63dpi and *tlr3* mutant hearts (n = 3 – 5 per time point) at 4dpi, 30dpi, 63dpi, 100dpi and 140dpi. Fibrotic scar clearance was assessed by measuring the fibrotic area as compared to the whole ventricular area using the ImageJ plug-in Fiji (National Institutes of Health, Bethesda, Maryland, USA) and calculating the relative amount affected by fibrosis. Regeneration efficiency analysis was performed by comparing the relative amount of fibrosis at 63dpi to the mean relative area injured at 4dpi in *tlr3* mutants and WT, respectively. On both analyses, sections with the largest relative amount injured compared to the whole ventricular area were measured.

### Immunofluorescence

*tlr3* mutant and WT hearts (n = 3 per group) were isolated at 6hpi as described above. Tissue sections were deparaffinized in xylene twice for 10’ and rehydrated for 5’ in decreasing concentrations of Isopropanol (100%, 96%, 70%, diH_2_O). Antigen retrieval was performed by incubation in Sodium-Citrate buffer (100mM Sodium Citrate, 0.05% Tween 20, pH 6.0) for 20’ at 60°C. Sections were then washed in flowing tap water for 10’ and in Dulbecco’s phosphate buffered saline + CaCl_2_ + MgCl_2_ (Gibco, DPBS). Sections were blocked in 10% normal goat serum (NGS, Agilent Dako), 2% bovine serum albumin (BSA, Albumin Fraktion V, NZ-origin, Carl-Roth), 1x DPBS (+calcium, +magnesium, Gibco) for 30’ at room temperature and washed three times with DPBS (+MgCl_2_, +CaCl_2_). Sections were incubated with either Dylight 594-conjugated isolectin B4 (IB4-594, Vector Laboratories, Burlingame, CA) for macrophage staining or anti-Mpx primary antibody (Genetex, San Antonio, TX) for neutrophil staining, at 1:200 dilution in 2% BSA/DPBS overnight at 4°C. The next day, sections were washed three times and anti-Mpx subjected tissue sections incubated with secondary antibody anti-rabbit AlexaFluor 488 (AF488, anti-rabbit goat polyclonal antibody, ab150077, Abcam) at 1:200 dilution in 2% BSA/PBS for 30’ in the dark at room temperature. IB4-594 subjected tissue sections were kept in DPBS (+MgCl_2_, +CaCl_2_) in the dark while anti-Mpx subjected sections were incubated with secondary antibody. Sections were washed and nuclei stained using DAPI (10mg/ml; Invitrogen, Life Technologies Corp., Carlsbad, CA) diluted at 1:1000 in 2% BSA/PBS for 90 – 120 seconds in the dark. At last, sections were washed twice, embedded in Prolong Diamond Antifade Mountant (Invitrogen, Life Technologies Corp., Carlsbad, CA) and covered with 15mm diameter high precision coverslips (Marienfeld Laboratory Glassware). Imaging was performed using the Leica Confocal Microscope SP8 gSTED (laser scanning confocal microscope, tandem scanner, gatedSTED, pulsed WLL) and Leica imaging software (Leica, Wetzlar, Germany) and images were processed using Fiji (National Institutes of Health, Bethesda, Maryland, USA). Injured ventricles were identified by disrupted cardiac muscle tissue and high abundance of nuclei. Injured ventricular area and abundance of neutrophils and macrophages were quantified manually using ImageJ/Fiji.

### Reverse transcriptase PCR and quantitative RT-PCR

Ventricular isolation was performed, and the BA removed afterwards. Whole ventricles from cryoinjured (*tlr3* mutants and WT) zebrafish were isolated at 6 hpi & 1 dpi (n = 3; 2 – 3 ventricles per sample). Sham treated ventricles were isolated 1 dps (n = 3; 2 – 3 ventricles per sample). Ventricles were snap-frozen in liquid nitrogen and stored in 1.5 ml microcentrifuge tubes at -80°C. For RNA-isolation, the Monarch® Total RNA Miniprep Kit (New England Bio Labs, Ipswich, MA) was used. Prior to RNA isolation, tissues were mechanically homogenized in 1.5 ml microcentrifuge tubes filled with 100 μl 1x DNA/RNA Protection reagent using a microtube pestle (VWR). Upon tissue homogenization, RNA isolation was performed following the Monarch® Total RNA Miniprep Kit instruction manual. Total RNA was eluted in 50 µl of nuclease free H_2_O (New England BioLabs). Subsequent cDNA generation was set up using the LunaScript® RT SuperMix Kit and its corresponding manual. qPCR reactions were set up using the Luna® Universal qPCR Master Mix according to the kit’s instructions manual. The following primers were used for the qPCR reaction concerning their respective genes. For the genes *il6st* (fwd: 5’-TCCTGAGCGTCTTCACCATA-3’, rvs: 5’- GCGGCCATAACAGCTTCTT-3’)*, jak1* (fwd: 5’-AAACACATCGCCCTGCTCTA-3’, rvs: 5’-AAAGGGCCGTACTGAACAAA-3’) and *stat3* (fwd: 5’-GGACTTCCCGGACAGTGAG-3’, rvs: 5’-ATCGCTTGTGTTGCCAGAG-3’), primers were designed as in Fang et al^22^. Primers for *csf1ra* (GCCCACATCCCATAATGCCT, CTCGCAACAGGCTTCGTGTA-3’)*, mpeg1* (fwd: 5’-CACAGAAAACCAGCGCATGAA-3’, rvs: 5’-CAGATGGTTACGGACTTGAACCC-3’) were generated using the NCBI Primer-Blast tool using default settings except for: Template (NCBI Gene ID), PCR product size (75 – 200 bp), Exon junction span (Primer must span an Exon-Exon junction), Primer Pair Specificity checking with Exclusion Organism (*Danio rerio).* As a reference gene, *eukaryotic elongation factor 1 alpha* (*eef1a1,* fwd: 5’-TCTCTACCTACCCTCCTCTTGGTC-3’, rvs: 5’- TTGGTCTTGGCAGCCTTCTGTG-3’) was used. Primer oligonucleotides were generated at Microsynth AG, Balgach, Switzerland. qPCR analysis was performed using Applied Biosystems 7500 Real Time PCR Systems and corresponding Applied Biosystems 7500 Software v2.0.5. SYBR® Green I dye reagent was used. Real time PCR was set up as follows: Initial Denaturation: 95°C / 1’; 40x cycles: Denaturation: 95°C / 15’’, Primer Annealing & Elongation: 60°C / 30’’; Exit Cycles; Denaturation: 95°C / 15’’, Annealing & Elongation: 60°C / 1’; Melt Curve Measurement: 60°C to 95°C / +1°C every 30’’; Final Denaturation: 95°C / 15’’, Final Annealing & Elongation: 60°C / 1’. Specific gene expression was normalized to the reference gene *eef1a1* given by the formula 2^-ΔCt^. The result for the Fold Change was calculated by the 2^-ΔΔCt^ method, normalized to corresponding sham treated ventricles. The mean Ct values were calculated from double determinations and samples were considered negative if the Ct values exceeded 35. Differences in Fold Change were considered as statistically significant at adjusted p-value < 5 x 10^-2^, assessed via unpaired t-test analysis assuming all measurements were sampled from populations with equal variance using GraphPad Prism (v8.4.3).

### RNA sequencing

Ventricular isolation was performed, and the BA removed afterwards. Whole ventricles from cryoinjured (*tlr3* mutants and WT) zebrafish were isolated at 1dpi & 4dpi (n = 2; 2 – 3 ventricles per sample). Sham treated ventricles were isolated 1dps (n = 2; 2 – 3 ventricles per sample). Ventricles were washed twice in 1x DPBS and transferred to 2mL screw cap tubes containing ∼20 sterile glass beads and submerged in 1mL TriReagent (Molecular Research Center, Cincinnati, Ohio, USA). Using a Tissue Homogenizer (Precellys Evolution Homogenizer), ventricles were mechanically disrupted (2x 5000rpm for 15 seconds, pause for 15 seconds in between) and incubated at room temperature for 5’. Then, 200µl chloroform (Carl Roth) was added and samples vortexed for 15 seconds. Upon further incubation for 2’ at room temperature, samples were centrifuged at 12 000g / 4°C / 15’. The upper, aqueous phase was transferred in 1.5mL microcentrifuge tubes and 1:1 volume of isopropanol (100%) was added (∼500µl). Using a pipette, samples were mixed and incubated for 30’ at -20°C and then for 10’ at room temperature. Afterwards, samples were centrifuged at maximum speed (∼20 000g) / 4°C / 30’ and the supernatant removed. The pellet was washed in 1mL 70% Ethanol (EtOH) and centrifuged at maximum speed / 4°C / 10’. Washing and subsequent centrifugation was repeated once. Supernatant was removed and the pellet air-dried under a laminar flow hood for 5 – 10’, until no EtOH residues were visible. The pellet was dissolved in 15µl nuclease free H_2_O and resuspended by pipetting. RNA was then subjected to DNase treatment by addition of 2µl DNase I RNAse free (Thermo Scientific), 2.5µl DNase buffer with MgCl_2_ (Thermo Scientific) and 5.5µl nuclease free H_2_O for 30’ at 37°C. Then, 2.5µl of 50mM EDTA (Thermo Scientific) was added, and samples further incubated for 5’ at 65°C. Afterwards, in that order, 160µl nuclease free H_2_O, 20µl ammonium acetate (NH_4_OAc, 5M) and 600µl 100% EtOH were added and mixed by pipetting. Samples were incubated for 60’ at -80°C and then centrifuged for 30’ at 20 000g / 4°C. Supernatant was removed, the pellet washed in 1mL 75% EtOH and samples centrifuged at maximum speed / 4°C / 10’. Washing was repeated twice. Supernatant was removed and the pellet was air-dried under a laminar flow hood, until no EtOH residues were visible (∼10’). The pellet was resuspended in 15µl nuclease free H_2_O and mixed by pipetting. RNA concentration and purity were measured with a NanoDrop 2000 Spectrophotometer (Thermo Scientific). RNA was submitted to Novogene (Cambridge, United Kingdom) for subsequent processing and sequencing. RNA quality was validated through Bioanalyzer 2100 system (Agilent Technologies). mRNA libraries were prepared via poly A enrichment library preparation. Bulk sequencing of every sample was performed using the NovaSeq 6000 sequencing system (Illumina). Sequencing resulted in > 20 million read pairs of 150bp paired-end reads per sample.

### RNAseq data preprocessing and differential expression analysis

Raw FastQ files were processed using the nf-core framework (REF). After UMI extraction, trimming of the adapters and low-quality sequences, the reads were mapped to the most recent version of the zebrafish genome (GRCz11) with STAR^40^. Transcripts were then quantified with Salmon^39^ using the zebrafish transcriptome annotation V4.3.2 from Lawson et al^40^. Raw counts and transcripts per million (TPM) were used for subsequent downstream analysis. Using the R package DESeq2, ver. 1.38.0^43^, DESeqDataSets were generated using the raw counts matrices as inputs. Gene filtering was performed, omitting genes with a summed total of >= 10 counts throughout all samples combined. Datasets were further normalized using the DESeq median of ratios method and log2-transformed. Principal Component Analysis (PCA) was performed on transformed data sets. Genes with absolute principal component 1 (PC1) loadings >= 0.02 were viewed as contamination from tissue other than heart tissue and excluded from downstream analysis. Differentially expressed genes (DEGs) were identified through DESeq2 analysis. Significant DEGs were classified with adjusted p-value < 5 x 10^-2^. Distribution of DEGs throughout different treatments and time points were displayed in a Venn diagram using Venny ver. 2.1.^41^. For heatmap generation, logarithmic values by the base of 10 (log_10_) of the normalized counts were calculated and used as inputs. Collagen subtypes and secreted factors were assembled from the zebrafish matrisome^15^ and aligned to the log_10_ normalized counts matrix. Heatmaps were generated using the R package pheatmap ver. 1.0.12^42^. Distinct dynamics of gene expression over a time axis of four days after inducing myocardial infarction were assessed and assigned to corresponding gene ontologies using the Java application of STEM (Short Time-series Expression Miner)^25^.

### Gene Set Enrichment Analysis

Raw counts were normalized through DESeq2 and normalized counts were then used for gene set enrichment analysis (GSEA)^43^. The java desktop application provided by the Broad Institute (University of Calilfornia San Diego, La Jolla, California, USA) was used to perform GSEA. Subsets of gene sets were extracted from the Molecular Signatures Database (MSigDB) using R. Hallmark gene sets (MSigDB collection = H), curated gene gets (MSigDB collection = C2), ontology gene sets (MSigDB collection = C5) and cell type signature gene sets (MSigDB collection = C8), assembled to the zebrafish genome, were analysed. To assess ventricular abundance of immune cells through distinct molecular patterns as well as patterns of cardiomyocyte dedifferentiation to a progenitor-like state, validated gene sets generated by single-cell transcriptional profiles in human skeletal muscle (as part of the MSigDB collection C8)^29^ were used. GSEAs were ran with default settings except for: Permutation type = gene_set; Metric for ranking genes = log2_Ratio_of_Classes; Max size: exclude larger sets = 1000. To evaluate expression differences throughout the whole dataset, all groups of WTs and all groups of *tlr3* mutants were compared within each other. Furthermore, gene set enrichment scores in WT 1dpi vs. *tlr3^-/-^* 1dpi and WT 4dpi vs. *tlr3^-/-^* 4dpi were assessed. To examine potential artifacts created by differing baseline expression dynamics between WT and *tlr3* mutants, GSEA was performed between WT Sham and *tlr3^-/-^*Sham. All GSEA results displayed did not show any enrichment when comparing WT to *tlr3* mutant sham treated controls, ruling out gene set enrichment due to *tlr3* knock-out without cryoinjury (as seen in Suppl. Table 1). GSEA was performed to display up- and downregulated gene set regulation patterns in WTs compared to *tlr3* mutants at certain time points directly and confirmed by showing the same pattern when running the GSEA comparing cryoinjured samples to their respective sham treated controls. Gene sets were classified as significantly differentially regulated with a False Discovery Rate (FDR) < 5 x 10^-2^.

### Cell Type Deconvolution

RNA deconvolution methods can infer cell fractions from bulk RNA-seq data leveraging pre-built genomic signatures, i.e. genomic fingerprints of a set of cell types, usually immune subsets. To infer immune cell fractions from RNA-seq data generated from the isolated ventricles, the quanTIseq pipeline was used^44^. This method was developed to deconvolve human RNA-seq data. To run analysis on zebrafish ventricles, the most recent version of the zebrafish genome (GRCz11) was remapped, aligning every zebrafish gene to its correspondent human ortholog. This information was used to remap TPM zebrafish RNAseq data to human genes. Zebrafish genes that were not successfully remapped were kept in the dataset with their original gene names. Zebrafish genes that mapped to multiple human genes were assigned to all the corresponding human orthologues. For multiple zebrafish genes mapped to a single human gene, the mean TPM of all these zebrafish genes was calculated and then assigned to the human gene. Differences in immune cell abundance between cryoinjured ventricles of WTs and *tlr3* mutants were considered as statistically significant at adjusted p-value < 5 x 10^-2^ assessed via unpaired t-test analysis, assuming all measurements were sampled from populations with the same scatter using GraphPad Prism (v8.4.3).

### Cell Culture

Wild-type and *TLR3^−/−^* human dermal fibroblasts were kindly provided by Jean-Laurent Casanova (Rockefeller University, New York, NY). Human dermal fibroblasts (hDFBs) were cultured and maintained in DMEM with 10% FCS and 1% penicillin/streptomycin/glutamine at 37°C and 5% CO_2_.

### Migration Assay

hDFBs (WTs and TLR3 mutants) were seeded in 6-Well Plates at 1.5 x 10^5^ cells per Well and starved for 24h in serum-free RPMI-1640 with 1% penicillin/streptomycin/glutamine at 37°C and 5% CO_2_. To stimulate TLR3 activation, hDFBs were then incubated with Polyinosinic:polycytidylic acid (Poly(I:C); Invivogen, San Diego, CA) in serum-free RPMI- 1640 at a concentration of 20µg/ml for 1h. Poly(I:C) was removed by washing the cells twice with PBS. Treated cells were then incubated in serum free RPMI-1640 for 24h and the media sterile-filtered. Control wells were incubated with the same media, but without Poly(I:C).

For migration assay applications, the QCM Chemotaxis 5µm 96-Well Cell Migration Assay Kit (Merck, Sigma-Aldrich, Darmstadt, Germany; Cat. No. ECM512). Human peripheral blood mononuclear cells (PBMCs) of three independent participants were isolated according to Cui et al^45^. All human experiments were approved by the ethics committee of the Medical University of Innsbruck (Ref. Nr.: 1262/2023). Filtered media from stimulated hDFBs was added to the wells of the feeder tray. PBMCs were loaded onto cell migration chambers at 2 x 10^5^ cells per chamber in serum-free RPMI-1640 with 1% penicillin/streptomycin/glutamine and put onto the feeder tray. For every treatment group and genotype, triplicates were set up comprising of one sample of PBMCs of each donor. PBMCs were incubated in treated media from hDFBs for 6 hours at 37°C and 5% CO_2_. Migrated PBMCs were further processed according to the manufacturers protocol and stained with CyQuant GR Dye diluted at 1/75 (Merck, Sigma-Aldrich, Cat. No. ECM512). Fluorescence was measured at 480 ± 15 nm (excitation, 480 ± 15 nm; dichroic mirror, auto, 500.8 nm; emission, 520 ± 12 nm; top excitation and detection) using CLARIOstar Plus microplate reader (BMG Labtech GmbH, Ortenberg, Germany) in 96-well cell culture microplates (Greiner, Kremsmünster, Austria; Ref.: 655090). A regression line was generated with increasing cell numbers (0, 100, 200, 500, 1000, 2000, 5000, 10000) to calculate the number of cells migrated through the migration chambers.

### Flow Cytometry

Isolated human PBMCs before and after migration were analyzed via Flow Cytometry. Cells were washed with 2% BSA in PBS and then incubated with APC-conjugated CD14 antibody (Invitrogen, eBioscience, Life Technologies Corp., Carlsbad, CA, Ref.: 17-0149-42), FITC-conjugated CD11b antibody (BD Biosciences, BD Pharmigen, Cat. No.: 562793) and 7-AAD (BioLegend, San Diego, CA, Cat. No.: 420403) for 30’ at room temperature in the dark at 1/100 dilution in 2% BSA in PBS. Cells were washed once more, resuspended in 2% BSA in PBS and 5 x 10^5^ events measured per sample using a LSRFortessa Flow Cytometer and FlowJo v10.9.0 (BD Biosciences).

### Olink

Proximal extension assays (Olink®) were used to compare protein levels released from WT and *TLR3* mutant hDFBs into the cell culture media upon Poly(I:C) (20µg/ml) stimulation for 24h at 37°C and 5% CO_2_. Relative expression levels of 92 proteins were determined using the Target 96 Inflammation Panel. Statistical significance was evaluated through Welch two-sided two sample t-test.

## Statistical Analysis

Data are presented as mean ± SEM for continuous variables, absolute numbers, and percentages for categorical variables. The confidence interval was set at 95%, therefore a p-value was deemed as 0.05. As appropriate, comparisons between groups were performed for continuous variables with unpaired, two-sided t-tests, corrected for multiple comparisons using the Holm-Sidak method. Data documentation and statistical analysis were performed using GraphPad Prism (v8.4.3) and RStudio Version 4.2.2 (RStudio Team, Boston, USA).

## Ethics

All human research participants included have provided consent to be included in this study. Every experiment including human participants has been approved by the ethics committee of the Medical University of Innsbruck (Ref. Nr.: 1262/2023). All experiments including the use of laboratory animals have been approved by the federal ministry of education, science, and research of Austria (Ref. Nr.: 2020-0.345.504).

## Data access and analysis statement

Manuel Fiegl has full access to all the data in the study and takes responsibility for its integrity and the data analysis.

## Results

### *tlr3* deletion alters survival and physical performance upon myocardial injury

To assess the effects of *tlr3* deficiency in cardiac repair, survival of zebrafish after myocardial cryoinjury was evaluated until 10 days post myocardial infarction (dpi). No deaths occurred beyond 10dpi. Survival analysis revealed an increased death threat in *tlr3* homozygous mutants subjected to cryoinjury as compared to WT controls, while there was no difference in survival of either group in sham treated zebrafish (**Fig. 1a**). These findings suggest inability of *tlr3* mutants to stabilize the ventricle upon cryoinjury, resulting in death. To evaluate potential impairment of cardiac function after injury in *tlr3* mutants, behaviour and physical performance were analysed. Fish react to oxygen deprivation by decreasing their distance to the water surface (DTS) and gasping for air, as the surface area provides the highest oxygen levels in the tank^24^. DTS of WT and *tlr3* mutants were evaluated until four weeks post cryoinjury treatment. DTS was decreased in *tlr3^-/-^* fish swarms, swimming closer to the water surface compared to WTs during the 4 weeks of investigation (**Fig. 1b,c; Suppl. Fig. 1**). Furthermore, *tlr3^-/-^* fish gasped for air more frequently than WTs (**Suppl. Video 1-3**). These results suggest increased oxygen deprivation in *tlr3^-/-^* treated fish upon myocardial injury, resulting in decreased DTS.

**Figure 1:**
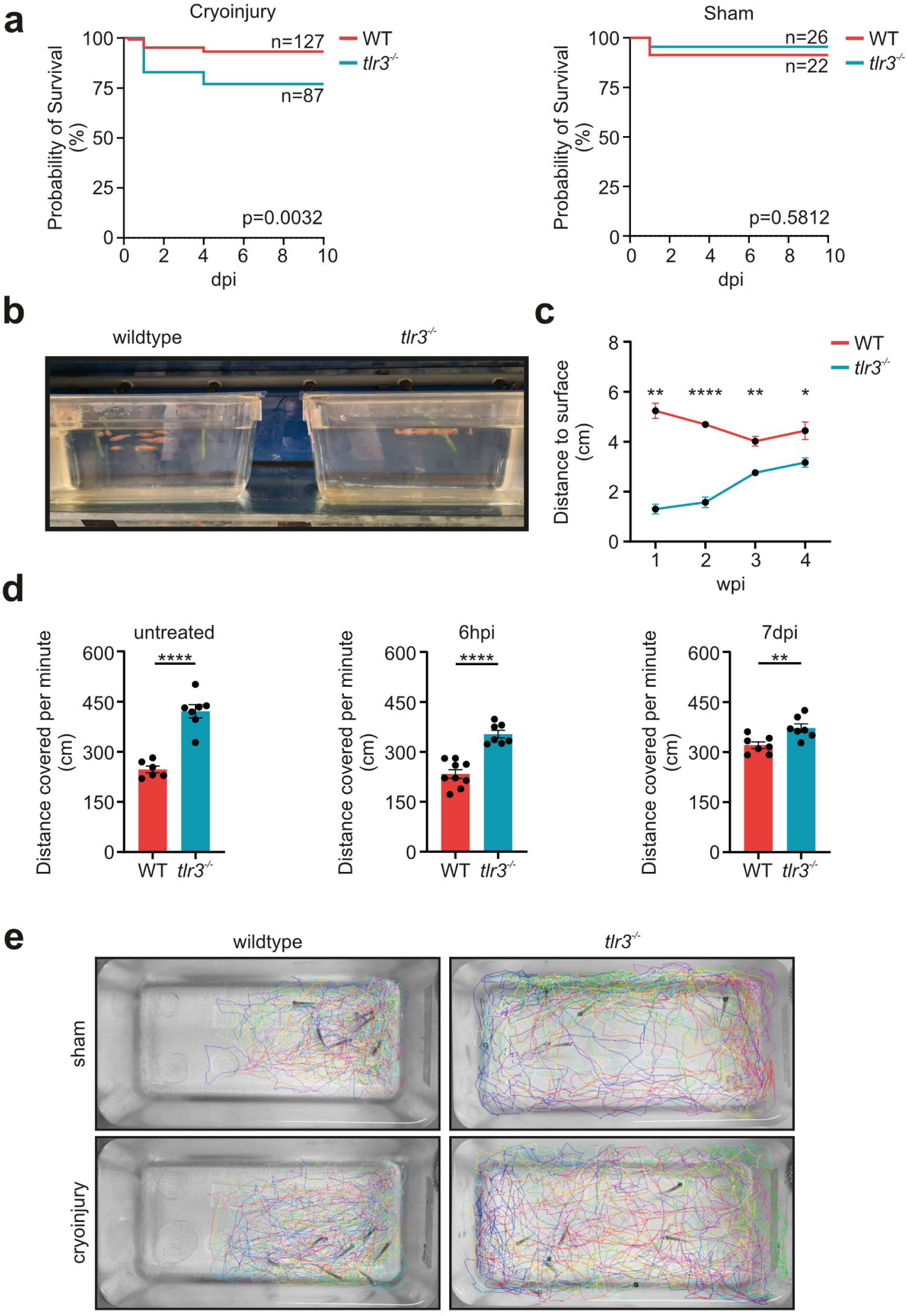
*tlr3* deficiency diminishes zebrafish survivability and alters movement behaviour upon cryoinjury. **a** Kaplan-Meier-Curve displaying survival probability of zebrafish upon subjection to cryoinjury and sham treatment in per cent. Mantel-Cox Log-rank p-test displays significance levels and results were viewed as statistically significant at p-value < 5 x 10^-2^. **b** Representative image taken for DTS evaluation. **c** DTS measures of *tlr3* mutants and WTs upon cryoinjury subjection. Curves displayed were generated calculating the mean DTS of all daily mean values per week. n = 8 – 11 fish were kept in every tank. Images were taken randomly during daytime n = 3 – 5 times per week. **d** Distance covered per minute measured by filming tanks containing n = 6 – 9 zebrafish subjected to respective treatments. Videos were taken randomly during daytime and untreated fish data was assessed 14 days upon putting them in their respective tanks. **e** Movement tracking of zebrafish for 60 seconds at 7 days post cryoinjury and sham treatment. Colours are corresponding to n = 7 fish per tank. **c, d** Data represents means ± SEM. * = adj. p-value < 5 x 10^-2^, ** = adj. p-value < 10^-2^, *** = adj. p-value < 10^-3^, **** = adj. p-value < 10^-4^.

Given these findings, we investigated whether *tlr3* knockout results in altered swimming activity upon myocardial infarction induction. *tlr3* mutants swam more actively in the tank than their WT counterparts, covering more distance at 6 hours post myocardial infarction induction (hpi), 7 dpi and 14 days after putting non-injured fish in mini swarms. The increased movement was found in cryoinjured and non-injured groups (**Fig. 1d**).

As *tlr3* mutant fish responded to a myocardial infarction with altered tank distributions in the z-axis, we further analysed tank occupation on a planar view. Wildtypes exhibited swarm like swimming behaviour close to one and another in the tank, ultimately occupying about two thirds of the tank area after a minute. *tlr3* mutants displayed a more heterogenous distribution, occupying the whole tank area after being subjected to both sham treatment and cryoinjury (**Fig.1e).** These findings show altered movement behaviour in *tlr3* mutants.

### Impaired myocardial scar remodelling in *tlr3* mutants

*tlr3* is a central mediator of cardiac regeneration in neonatal mice and play a pivotal role in the regeneration of other tissues such as the spinal cord^19,20^. Hence, we aimed to assess the ability of *tlr3* knock out zebrafish to resolve fibrotic scar tissue after myocardial infarction. Therefore, myocardial injury was induced, and zebrafish hearts histologically examined over a period of 140 dpi. There was no significant difference in injury severity between the groups at 4dpi(**Fig. 2a, c**). To evaluate scar remodelling of *tlr3* mutant zebrafish two months after myocardial infarction induction, the resolution of the relative injured area was compared at 4 dpi, 30 dpi 63 dpi. WT fish cleared 81% (± 6% SEM) of their initially formed fibrosis, whereas *tlr3* mutants resolved only 43% (± 10% SEM) (**Fig. 2d**). Moreover, *tlr3* mutants failed to significantly resolve fibrosis until 63 dpi, while WTs reduced fibrosis below 10% of the ventricular area at that time point **(Fig. 2a, c**). Given the fact that WT zebrafish resolve a cryoinjury induced cardiac fibrosis entirely at ∼100 dpi^2,6^, we evaluated the scar size of *tlr3* mutants beyond that time point. At 140 dpi, mutant ventricles still displayed similar levels of fibrosis as at 63 dpi, whereas cardiac tissue of WTs had entirely regenerated (**Fig. 2b**). These findings suggest that *tlr3* signalling is a central driver in the resolution of myocardial fibrosis in zebrafish.

**Figure 2:**
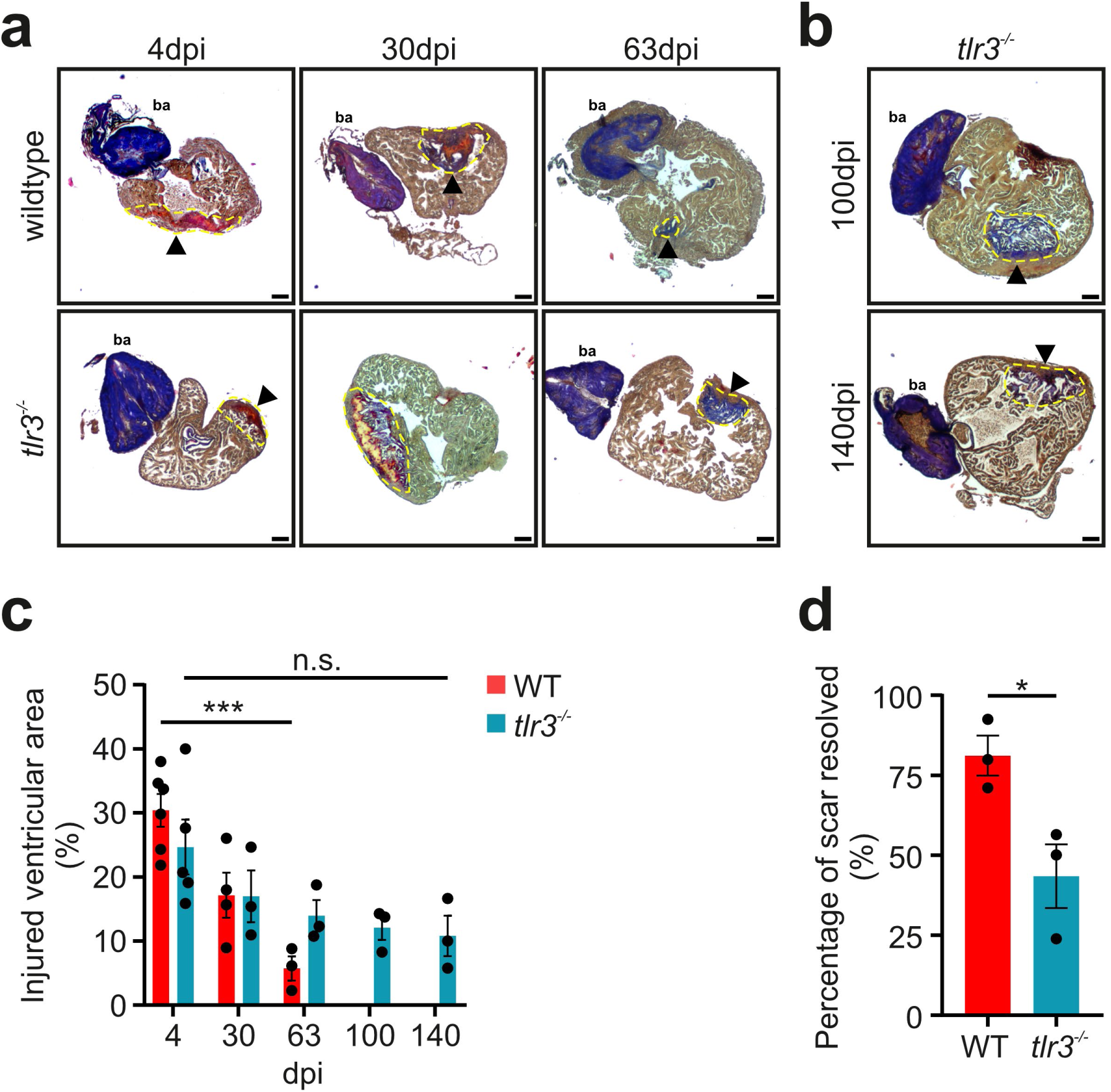
*tlr3* mutants display diminished scar resolution after cardiac injury. **a, b** AFOG stained histological sections of cryoinjured ventricles (section thickness = 5µm). Fibrosis is stained in blue (collagen) and red (fibrin), whereas healthy cardiac muscle is coloured yellow/orange. The bulbus anteriosus (ba) is located anterior to the ventricle, stained in blue. Injured area is highlighted in yellow dashed lines. Arrowheads indicate the injured ventricular area. scale = 100µm. **b** Sections of *tlr3* mutants at 100dpi and 140dpi. **c** Relative amounts of the cardiac ventricle subjected to fibrosis. Injured areas were measured and compared to whole ventricular areas. n = 3 – 6. dpi = day post myocardial infarction. **d** Regeneration efficiency of the ventricle as measured by calculating the relative amount of fibrosis resolved throughout 2 months by comparing extents of fibrosis at 63 dpi to the mean relative area injured at 4 dpi. n = 3 – 6. **c, d** Data represents means ± SEM. * = adj. p-value < 5 x 10^-2^, ** = adj. p-value < 10^-2^, *** = adj. p-value < 10^-3^, **** = adj. p-value < 10^-4^.

### Reduced myocyte dedifferentiation signatures in *tlr3^-/-^*

To identify molecular mechanisms underlying reduced survival in *tlr3* mutants, fibrosis clearance and cardiac repair, bulk RNA sequencing (RNAseq) was performed at 1dpi and 4dpi and compared to sham treated controls. Principal component analysis (PCA) revealed uncommonly high heterogeneity in gene expression between sham treated replicates in WT and *tlr3^-/-^* fish along the first principal component (PC1) (**Suppl. Fig. 2a**). The genes associated with PC1 (i.e. 286 genes with absolute PC1 loadings ≥ 0.02) were expressed solely in sham treated outlier samples. As these genes are not canonically expressed in cardiac muscle (**Suppl. Fig. 2b, c; Suppl. Table 1**) and might, thus, be due to tissue contamination, we excluded them from downstream RNAseq analysis. This filtering resulted in a PCA plot separating WTs and *tlr3* mutants (**Fig. 3a**). Differential gene expression analysis revealed statistically significant differences in the expression of numerous genes between WTs and *tlr3* mutants (**Fig. 3b**).

**Figure 3:**
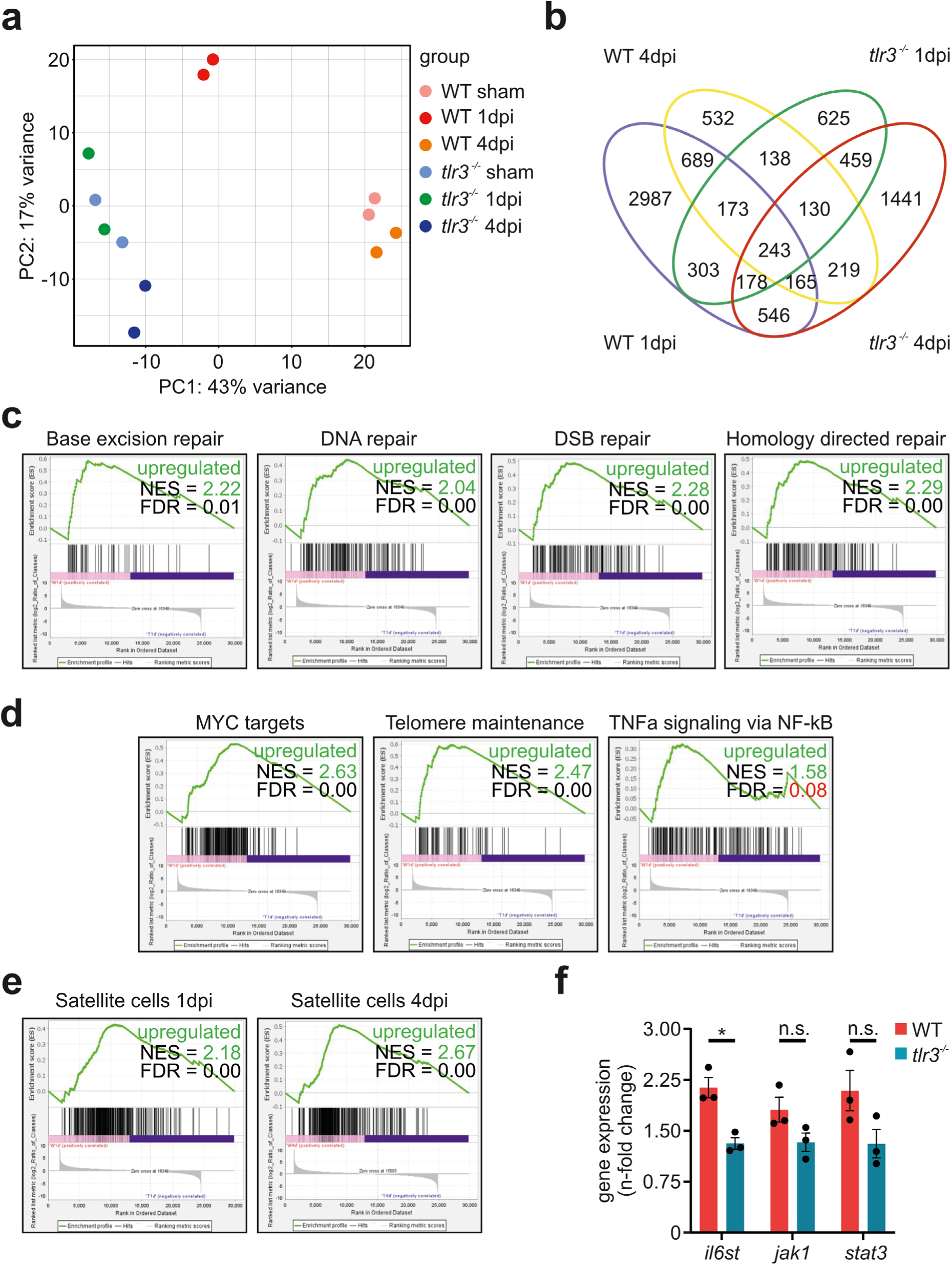
*tlr3* mutants undergo diminished cardiomyocyte dedifferentiation. **a** PCA-plot depicting transcriptome heterogeneity between WT and *tlr3* mutant ventricles subjected to cryoinjury or sham treatment. n = 2. **b** Venn diagram displaying the distribution of significant DEGs (adj. p-value < 5 x 10^-2^) between WTs and *tlr3* mutants at given time points. **c** GSEA enrichment plots at 1dpi for DNA-repair mechanisms of validated gene sets of given REACTOME, Wikipathways and Hallmark gene sets. DSB = double strand break. NES = normalized enrichment score. FDR = false discovery rate. Positive NES values display enrichment in WTs compared to *tlr3* mutants, as highlighted in green. **d** Enrichment plots at 1dpi for gene sets of given REACTOME and Hallmark gene sets. FDRs > 5 x 10^-2^ are highlighted in red. **e** Enrichment plots of the gene set skeletal muscle satellite cells, generated by Rubenstein et al^30^. **f** quantitative real-time PCR displaying 2^-ΔΔCt^ values as Fold Change, normalized to *eukaryotic translation elongation factor 1 a* (*eef1a1*) and respective sham treated samples. *il6st = interleukin 6 cytokine family signal transducer. jak1 = janus kinase1. stat3 = signal transducer and activator of transcription 3.* Data represents means ± SEM. ns = not significant, * = adj. p-value < 5 x 10^-2^.

Enriched toll-like receptor signalling in WTs compared to *tlr3^-/-^*was confirmed using STEM (Short Time-series Expression Miner^25^) (**Suppl. Fig. 2d**).

Using GSEA, we discovered upregulation of genes associated with several DNA-repair subtypes, such as Base Excision, Homology Directed and Double Strand Break repair in WTs at 1dpi compared to *tlr3* mutants (**Fig. 3c**). Protection of cardiomyocytes followed by subsequent dedifferentiation is pivotal for effective cardiac repair in zebrafish^2,4,18^. Upon cryoinjury, extensive tissue apoptosis results in the loss of cardiomyocytes^6^. A central mediator of apoptosis is the accumulation of DNA-damage due to unsuccessful repair^26^. Activation of DNA-repair mechanisms in adult mouse cardiomyocytes alleviates DNA-damage upon myocardial infarction induction, resulting in improved cardiac function with subsequent reduced cardiac tissue cell loss and increased myocyte proliferation^4^. When looking at the gain of proliferative capacity in cardiomyocytes, certain signalling cascades have been shown to be indispensable. Nf-κb, a transcription factor shared among different TLRs including Tlr3, drives dedifferentiation in cardiac myocytes^23^. In parallel, Nf-κb signalling leads to the expression of *tumor necrosis factor α* (*tnfα*), a cytokine expressed early after injury in macrophages migrating towards the injured ventricle^14^. We found enrichment of Tnfα signalling via Nf-κb in WT injured ventricles at 1dpi, compared to *tlr3* mutants. Furthermore, targets of Myc, which is implicated in cellular plasticity^27^, were enriched in WTs compared to *tlr3* mutants at 1dpi. Moreover, genes responsible for telomere maintenance, which are active only in stem-cells and cells at a progenitor-like state with increased proliferative capacity^28^, were upregulated (**Fig. 3d**). Another target of *tlr3* mediated *nf-κb* activation is the *il6/jak1/stat3* signalling cascade, driving cardiac muscle tissue proliferation and dedifferentiation in zebrafish. *Interleukin 6 cytokine family signal transducer (il6st)* binds *interleukin 6* (*il6*) and triggers signalling towards the intracellular domain of targeted cells^22^. *Il6st* expression was increased in in WTs, while *tlr3* mutants failed to upregulate *il6st* mRNA levels at 1dpi (**Fig. 3f**). Ultimately, genes specifically active in satellite cells, which resemble the stem cell niche of muscle cells, were enriched at 1dpi and 4dpi in WTs compared to *tlr3* mutants (**Fig. 3e**). Altogether, these findings indicate a central role of *tlr3* mediated signalling in finetuning molecular patterns of cardiomyocyte dedifferentiation and proliferation.

### Impaired immune cell migration and maturation in *tlr3^-/-^*

Myeloid immune cells, particularly macrophages and neutrophils, are key mediators of cardiac repair in zebrafish^2,14,30^. Attracted by cell debris, damage-associated molecular patterns (DAMPs) and cytokines, myeloid immune cells extensively infiltrate the injured ventricular area immediately upon myocardial infarction. Neutrophils are the first immune cells to migrate towards the injured ventricle, followed by macrophages^2,31^. Since TLR3 activation orchestrates innate immune system responses, we aimed to assess the macrophage and neutrophil immune cell abundance in the injured ventricle of *tlr3* mutant zebrafish. Immunofluorescent imaging revealed significantly lower numbers in neutrophils and macrophages in the injured ventricle of *tlr3* mutants at 6hpi, compared to WTs (**Fig. 4a-d**). These findings are further underlined by decreased expression of the macrophage marker *macrophage expressed gene 1* (*mpeg1*) in *tlr3* mutant ventricles (**Fig. 4e**). Notably, mutant hearts eventually reached similar expression levels of *mpeg1* as WTs at 1dpi, but these macrophages failed to polarize to their distinct phenotype normally present at this time point. Canonically, macrophages are divided into two groups. M1 macrophages are implicated in pro-fibrotic phenotypes, while M2 macrophages are mostly present in pro-regenerative phenotypes, such as resolution of inflammation and tissue repair^12,13^. Immune cell abundance measurements through cell type deconvolution revealed significantly reduced pro-regenerative M2 macrophages in *tlr3* mutant as compared to WT ventricles (**Fig. 4f**). In zebrafish, macrophages polarize towards a M2-like state through induction of *colony stimulating factor 1 receptor a* (*csf1ra*) expression at 1dpi, mediating the required fibrosis build-up in an organized manner early after cardiac injury^14^. Consistent with reduced M2 polarization, expression of *csf1ra* was strongly reduced in *tlr3* mutants (**Fig. 4g**). To further strengthen our findings regarding diminished immune cell abundance, we found significant enrichment in gene expression associated with the abundance of myeloid cells in WT ventricles compared to *tlr3* mutants using GSEA (**Fig. 4h**). These results suggest that disruption of *tlr3* signalling leads to diminished innate immune cell migration towards the injured ventricular area early after injury, resulting in diminished scar resolution, reduced cardiomyocyte dedifferentiation and impaired survival.

**Figure 4:**
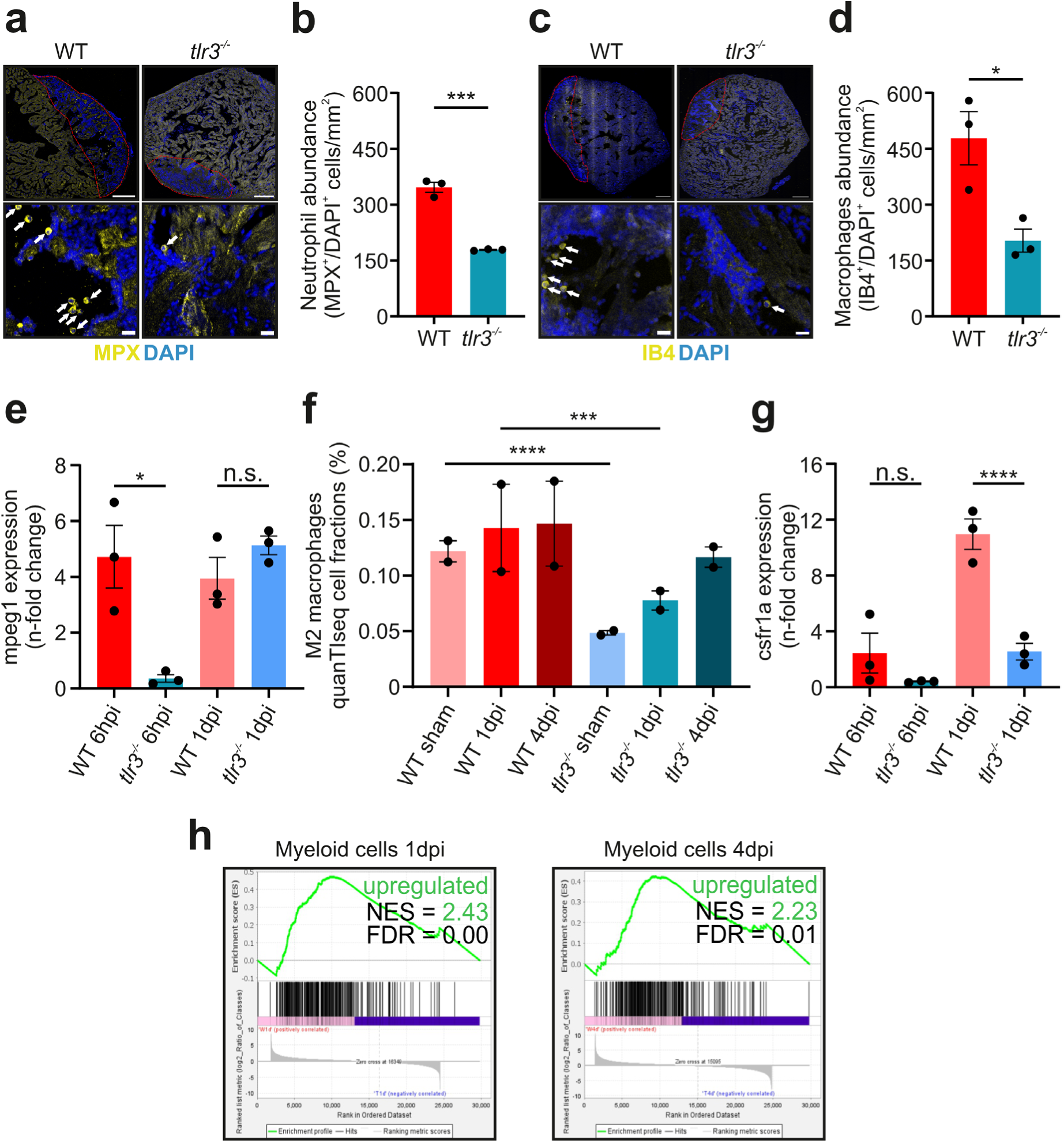
*tlr3* deficiency decreases myeloid immune cell recruitment to the injured heart. **a, c** Immunofluorescence images of sectioned ventricles subjected to cryoinjury. top: Imaging performed at 93x magnification, stitching single pictures afterwards. bottom: Representative 93x magnified images of injured areas displaying neutrophils (**a**) and macrophages (**c**). White arrows indicate counted immune cells. *myeloid specific peroxidase* (*mpx*) was stained in yellow to localize neutrophils (**a**) and macrophages were stained in yellow with isolectin B4 (IB4) (**c**). Nuclei are stained in blue using DAPI. Injured area is highlighted by red dashed lines. Nuclei surrounded by *mpx* or IB4 signal were counted as immune cells. top: scale = 100µm. bottom: scale = 10µm. **b, d** Immune cells abundant in the injured ventricular area counted through immunofluorescence imaging. **e, g** quantitative real-time PCR results, displaying Fold Change. *mpeg1* = *macrophage expressed gene 1. csf1ra* = *colony stimulating factor 2 receptor a.* **f** Cell type deconvolution using quanTIseq^31^. Data represent absolute values in per cent, predicting fractions of M2 macrophages abundant in ventricles used for RNAseq. **b, d, e, f, g** Data represents means ± SEM. * = adj. p-value < 5 x 10^-2^, ** = adj. p-value < 10^-2^, *** = adj. p-value < 10^-3^, **** = adj. p-value < 10^-4^. **h** GSEA enrichment plots for immune cell signatures of skeletal muscle myeloid cells at 1dpi and 4dpi. NES = normalized enrichment score. FDR = false discovery rate. Positive NES values display enriched gene sets in WTs compared to *tlr3* mutants, as displayed in green & gene sets were deemed significantly enriched at FDR < 5 x 10^-2^.

### ECM organization is dysregulated in *tlr3^-/-^* hearts

ECM constituents such as collagens stabilize the injured ventricle while also playing a central role in mechanical and biochemical interactions determining cell behaviour and function. Macrophages are the main mediators of the fibrotic response upon cardiac injury by inducing collagen deposition^14^. Since macrophage and neutrophil abundance is severely restricted in injured *tlr3* mutant ventricles, we expected *tlr3* mutant hearts to display altered complexity and composition of the fibrotic scar early after injury. Genes associated with collagen formation were enriched at 4dpi in *tlr3* mutants compared to WTs (**Fig. 5a**). 18 collagen genes were enriched in *tlr3^-/-^* infarcted ventricles. Of all collagen genes upregulated, half were enriched found in *tlr3^-/-^*exclusively, while the other half was as well upregulated in WT injured ventricles. (**Fig. 5b,c**). Every collagen gene enriched in both WT and mutant infarcted ventricles displayed higher expression levels in *tlr3* mutants (**Fig. 5c**). Since we found strong heterogeneity of collagen subtypes expressed between WT and *tlr3* mutants, we subsequently analysed whether differentially expressed collagen genes in *tlr3* mutants were implicated in phenotypes lacking fibrotic resolve. *Collagen type 11* is enriched in *chemokine receptor type 1* (*Ccr1*) homozygous mutant hearts of mice subjected to myocardial infarction. *Ccr1^-/-^* mouse ventricles display reduced neutrophil recruitment, resulting in stable collagen deposition and impaired cardiac repair^32^. *Collagen type 11* zebrafish orthologues *col11a1a* and *col11a1b* were both significantly enriched in *tlr3* mutant ventricles (**Fig. 5b,c**). Taken together, our findings suggest that the disrupted immune cell abundance causes disorganised fibrotic scar formation, ultimately leading to increased mortality, reduced cardiac function and incomplete cardiac repair.

**Figure 5:**
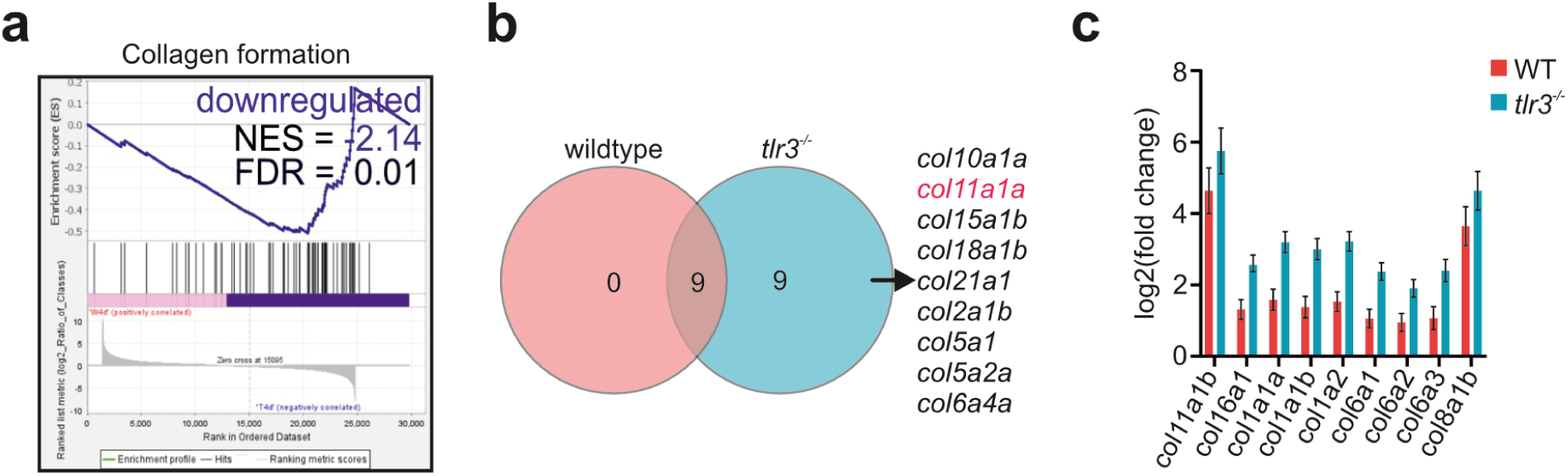
Enriched collagen formation and heterogeneity in *tlr3^-/-^* hearts. **a** Enrichment plots at 4dpi for REACTOME gene set collagen formation. NES = normalized enrichment score. FDR = false discovery rate. Negative NES values depict enrichment in *tlr3* mutants compared to WTs, as displayed in blue. Gene sets were deemed significant with FDR < 5 x 10^-2^. **b** Venn diagram showing distribution of significantly upregulated collagen genes between WT and *tlr3* mutant ventricles at 4dpi. **c** Expression of collagens upregulated in both groups. Data represents means ± SEM generated through DESeq2 analysis.

### Human fibroblast TLR3 activation increases chemotaxis

To determine whether the ability of activating TLR3 signalling directly contributes to increased immune cell migration, we assessed the ability of certain tissue resident cells to exert chemotactic stimulation. Cardiac fibroblasts are an essential cell type for extracellular matrix homeostasis and directly contribute to inflammatory cell infiltration through cytokine secretion upon myocardial infarction^33^. Hence, we expect fibroblasts to be the first major responders to tissue damage upon TLR3 activation. We stimulated human dermal fibroblasts (hDFBs) with the TLR3-agonist Polyinosinic:polycytidylic acid (Poly(I:C)) for 24 hours and analysed cytokine levels released into the supernatants. We found a substantial increase in chemotaxis factors stimulating monocyte, macrophage, and T-cell migration, such as *Monocyte Chemoattractant Protein 1* (*MCP-1*), *Monocyte Chemoattractant Protein 2* (*MCP-2*), *Monocyte Chemoattractant Protein* (*MCP-3*), as well as *C-X-C motif chemokine 10* (*CXCL10*). Additionally, besides the well-known induction of interferon-respnsive genes (circulation paper) we discovered an increase of *IL6* levels, a cytokine directly activated by TLR3 signalling and indispensable for zebrafish myocardial tissue regeneration^22,30^. We further found an increase in *Colony Stimulating Factor 1* (*CSF-1*) levels, which has been shown to drive macrophage maturation in early stages of zebrafish heart regeneration^14^ (**Fig. 6a,b**).

**Figure 6:**
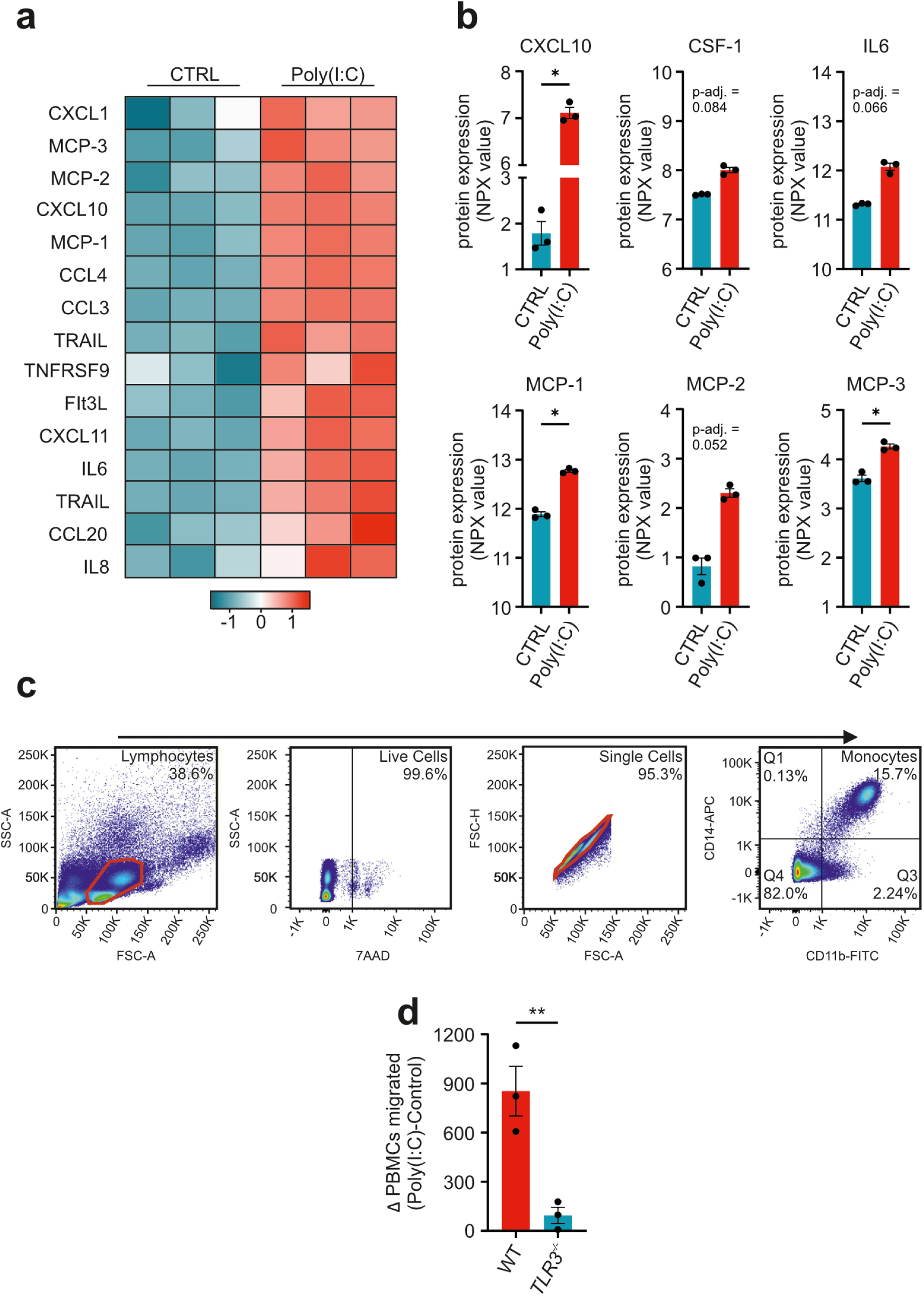
TLR3 signalling is crucial for immune cell migration. **a** Heatmap generated via the Olink® Statistical Analysis app depicting cytokines implicated in immune cell migration and maturation enriched in the media of Poly(I:C) treated WT hDFBs. **b** Relative levels of cytokine release into the media of Poly(I:C) treated hDFBs assessed via Olink® proximal extension assays. *MCP-1,2,3* = *Monocyte Chemotactic Protein 1,2,3. CSF-1* = *Colony Stimulating Factor 1. CXCL10* = *CXC Motif Chemokine Ligand 10. IL6 = Interleukin 6.* **c** Representative image of flow cytometry analysis of isolated human PBMCs. Live cells were assessed as 7-AAD negative. Mature Monocytes were viewed as double positive for CD14 and CD11b, after pre-gating for Lymphocytes. **d** Difference of cells migrated between PBMCs incubated with media from Poly(I:C) treated and untreated WT and *TLR3* mutant hDFBs. **b, d** Data represents means ± SEM. ns = not significant, * = adj. p-value < 5 x 10^-2^, ** = adj. p-value< 10^-2^.

Next, we aimed to elucidate the role of fibroblast TLR3 activation in monocyte and macrophage recruitment at a functional level. Therefore, we performed cell migration assays, assessing the number of human peripheral blood mononuclear cells (PBMCs) migrating towards a chemotactic source. Isolation of human PBMCs was confirmed using flow cytometry (**Suppl. Fig. 3**). PBMCs were incubated with supernatants from either WT of *TLR3* knock-out hDFBs treated with Poly(I:C) and the number of cells migrated through the 5µm pores compared to those of PBMCs incubated with supernatants from untreated hDFBs. After 6 hours of incubation, we found significantly enhanced migration of PBMCs towards supernatants deriving from PolyI:C stimulated fibroblasts. In contrast, supernatants from PolyI:C stimulated *TLR3* knock-out human fibroblasts did not alter PBMC migration rates, ruling out potential off- target effects of Poly(I:C) on PBMC migration (**Fig. 6d, Suppl. Fig. 4**). Taken together, these findings indicate that tissue resident TLR3 activation results in increased innate immune cell migration, potentially revealing a conserved mechanism of myocardial regeneration induction also in humans.

## Discussion

### Animal model of cardiac regeneration

Regenerative species like zebrafish retain latent ability to regenerate cardiac tissue, in contrast to non-regenerative species (e.g. humans). Against the decade-long perception of mammalian cardiac tissue having only little regenerative capacity, the finding that newborn mice can regenerate cardiac injury until seven days after birth led to suggesting that cardiac regeneration can be induced also in humans^3,5^.

### Innate Immunity and organized fibrosis establishment

*Tlr3* is an important mediator of innate tissue regeneration^19,20^. While *tlr3* undoubtedly plays a central role in mediating repair, no functional studies of *tlr3* knock-out mutant zebrafish in respect to cardiac injury resolution have yet been described. *tlr3* mutant zebrafish fail to recruit neutrophils and macrophages to the injured area of the heart. Both these cell types drive essential mechanisms of cardiac repair^2,10,11,13,14,34^. Different macrophage subtypes are responsible for the finetuning of the inflammatory responses, mediating formation of a fibrotic scar early after injury while later driving scar resolution and tissue regeneration^14^. Furthermore, neutrophils have been revealed to polarize highly plastic macrophages towards a pro-regenerative phenotype^35^. Therefore, we postulate that the reduced immune cell migration upon disrupted *tlr3* signalling leads to an altered inflammatory response in the injured tissue, affecting both scar build-up as well as resolution. Mutant fish seem to fail at establishing fibrosis in an organized manner necessary to stabilize the ventricle, resulting in impaired survival within the first ten days post injury.

### *Tlr3* signalling drives cardiomyocyte dedifferentiation

Besides organised fibrosis, regeneration is heavily dependent on the proliferative capacity of the affected tissue. Upon injury, zebrafish cardiomyocytes gain ability to proliferate through dedifferentiation to a progenitor-like state^2,18^. Cardiac muscle tissue protection by upregulation of DNA-repair mechanisms improves heart regeneration even in adult mice^4^. Here, we show upregulation of important DNA-repair patterns shortly after cryoinjury in WTs, but not in *tlr3* mutants. This is in line with our findings that *tlr3* knock-out leads to the downregulation of dedifferentiation markers implicated in processes such as telomere maintenance and the activity of target genes of the pluripotency marker *myc*. Strikingly, gene expression patterns specifically active in muscle stem cells were shown to be downregulated in *tlr3* mutants, further underlining our hypothesis of disrupted *tlr3* signalling leading to incomplete cardiomyocyte dedifferentiation.

### *Tlr3* mediated cardiac regeneration is evolutionary conserved

At last, we discovered a mechanism of human fibroblasts releasing specific cytokine signatures involved in both immune cell chemotaxis and maturation upon TLR3 activation. These cytokines resulted in the recruitment of PBMCs towards the chemotactic source, potentially unveiling on how ventricular resident fibroblasts migrate the necessary immune cells towards the injured area and facilitate the maturation towards their distinct phenotype, ultimately setting the starting point of the downstream regeneration cascade that eventually leads to complete scar resolution. Given the fact, that fibroblasts, directly contribute to inflammatory cell infiltration through cytokine secretion upon myocardial injury^33^, we identified fibroblasts as potential first responder cells able to improve cardiac regeneration upon TLR3 activation.

## Conclusion

Here, we provide novel evidence of *tlr3* being a key driver of cardiac regeneration in zebrafish. Survival of *tlr3* homozygous mutant zebrafish mutants was markedly impaired after myocardial injury compared to wild-types and mutant fish failed to complete scar resolution, mainly due to impaired neutrophil and macrophage migration to the injured ventricle and subsequent maturation. Mutant injured hearts too displayed increased heterogeneity in collagen gene expression. Ultimately, we postulate a potential conserved mechanism of *tlr3* activation in human fibroblasts to mediate immune cell migration towards the injured ventricular area and drive monocyte maturation. Taking all these findings into account, we demonstrate the crucial role of *tlr3* signalling in cardiac regeneration in a previously undescribed mutant model, providing further knowledge to the highly regulated processes of cardiac regeneration in zebrafish, one of the most relevant animal models in respect to heart regeneration. Our data reveal *tlr3* as a novel therapeutic target to potentially promote cardiac regeneration also in humans with ischemic cardiomyopathy.

## Acknowledgments

This work was supported by grants from the Austria Science fund (FWF) to Dr Gollmann-Tepeköylü (P 32821). This study is supported by VASCage – Research Center on Vascular Aging and Stroke. VASCage is a COMET Center within the Competence Centers for Excellent Technologies (COMET) program and funded by the Federal Ministry for Climate Action, Environment, Energy, Mobility, Innovation and Technology, the Federal Ministry of Labour and Economy, and the federal states of Tyrol, Salzburg, and Vienna. No use of artificial intelligence programs contributed to the compilation of the submitted manuscript in any way.

Each of the authors confirms that this manuscript has not been previously published and is not currently under consideration by any other journal. Additionally, every author has approved the contents of this paper and has agreed to the *JAHA* submission policies.

Each named author has substantially contributed to conducting the underlying research and drafting this manuscript. Additionally, to the best of our knowledge, the named authors have no conflict of interest, financial or otherwise.

## Abbreviations

Dpi: days post myocardial infarction
*tlr3*: *toll-like receptor 3*
WT: Wildtype
*col11*: *collagen type 11*
IHD: ischaemic heart disease
ICMP: ischaemic cardiomyopathy
SFPQ: *DNA repair- related molecule splicing factor, proline- and glutamine-rich*
DAMPs: danger associated molecular patterns
ECM: extracellular matrix
*plk1*: *polo-like kinase 1*
MI: myocardial infarction
*tlr*: *toll-like receptor*
dsRNA: double-stranded RNA
SWT: shock wave therapy
*myd88*: intracellular signaling adaptor myeloid differentiation primary response gene 88
*trif*: *TIR-domain-containing adaptor-inducing interferon-β*
IFN: type I interferons
*il6*: *interleukin- 6*
*irf*: *interferon regulatory factor*
*nf-κb*: *nuclear factor k-lightchain-enhancer of activated B cells*
PCA: principal component analysis
SEM: standard error of the mean
hpi: hours post myocardial infarction
*col11a*: *collagen type 11a*
gRNA: guide RNA
diH2O: distilled Water
bp: base pair
PFA: paraformaldehyde
BA: bulbus anteriosus
DTS: average distance to the water surface
NGS: normal goat serum
BSA: bovine serum albumin
DPBS: Dulbecco’s phosphate buffered saline
IB4-594: Dylight 594-conjugated isolectin B4
AlexaFluor 488: AF488
dps: days post sham
EtOH: ethanol
NH4OAC: ammonium acetate
tpm: transcripts per million
DEGs: differentially expressed genes
log10: logarithmic values by the base of 10
GSEA: gene set enrichment analysis
FDR: False Discovery Rate
wpi: week post injury
*mpeg1*: macrophage expressed gene 1
*mpx*: *myeloid specific peroxidase*
NES: normalized enrichment score
*il6st*: *interleukin 6 signalling transducer*
*jak1*: *janus kinase 1*
*stat3*: *signal transducer and activator of transcription 3*
*ccr1*: *chemokine receptor type 1*
hDFBs: human dermal fibroblasts
poly(I:C): polyinosinic:polycytidylic acid
*MCP-1*: *Monocyte Chemoattractant Protein 1*
*MCP-2*: *Monocyte Chemoattractant Protein 2*
*MCP-3*: *Monocyte Chemoattractant Protein 3*
*CXCL10*: *C-X-C motif chemokine 10*
*CSF-1*: *Colony Stimulating Factor 1*
*PBMCs*: *human peripheral blood mononuclear cells*
ICM: ischemic cardiomyopathy
SWT: shock wave therapy

## Data Availability

The data that support the findings of this study are available from the corresponding author upon reasonable request.

## Code Availability

The code used to generate findings of this study are available from the corresponding author upon reasonable request.

